# Promotion of RNF168-Mediated Nucleosomal H2A Ubiquitylation by Structurally-defined K63-Polyubiquitylated Linker Histone H1

**DOI:** 10.1101/2024.07.22.604500

**Authors:** Qiang Shi, Zhiheng Deng, Liying Zhang, Zebin Tong, Jia-Bin Li, Guo-Chao Chu, Huasong Ai, Lei Liu

## Abstract

The chemical synthesis of histones with homogeneous modifications is a potent approach for quantitatively deciphering the functional crosstalk between different post-translational modifications (PTMs). Here, we developed an expedient site-specific (poly)ubiquitylation strategy (CAEPL, Cysteine-Aminoethylation coupled with Enzymatic Protein Ligation), which integrates the Cys-aminoethylation reaction with the process of ubiquitin-activating enzyme UBA1-assisted native chemical ligation. Using this strategy, we successfully prepared monoubiquitylated and K63-linked di- and tri-ubiquitylated linker histone H1.0 proteins, which were incorporated into individual chromatosomes. Quantitative biochemical analysis of different RNF168 constructs on ubiquitylated chromatosomes with different ubiquitin lengths demonstrated that K63-linked polyubiquitylated H1.0 could directly stimulate RNF168 ubiquitylation activity by enhancing the affinity between RNF168 and chromatosome. Subsequent cryo-EM structural analysis of the RNF168/UbcH5c–Ub/H1.0–K63-Ub_3_ chromatosome complex revealed the potential recruitment orientation between RNF168 UDM1 domain and K63-linked ubiquitin chain on H1.0. Finally, we explored the impact of H1.0 ubiquitylation on RNF168 activity in the context of asymmetric H1.0–K63-Ub_3_ di-nucleosome substrate, revealing a comparable stimulation effect of both the inter- and intra-nucleosomal crosstalk. Overall, our study highlights the significance of access to structurally-defined polyubiquitylated H1.0 by CAEPL strategy, enabling in-depth mechanistic investigations of *in-trans* PTM crosstalk between linker histone H1.0 and core histone H2A ubiquitylation.

## INTRODUCTION

Cytotoxic DNA double-strand breaks (DSBs) trigger ubiquitylation signal cascades to recruit DNA repair factors to the damaged sites. Early in the DSB response, chromatin ubiquitylation is initiated by the sequential actions of two pivotal E3 ubiquitin ligases RNF8 and RNF168.^1-4^ It is reported that RNF8 catalyzes K63-linked polyubiquitylation of linker histone H1, which in turn recruits RNF168 to DSB sites.^5-7^ RNF168 then targets the N-terminal Lys 13/15 residues of nucleosomal H2A for ubiquitylation (H2AK13/15Ub), which subsequently facilitates the recruitment of downstream DNA damage repair factors like 53BP1 in the nonhomologous end-joining (NHEJ) pathway and BRCA1/BARD1 in the homologous recombination (HR) pathway.^8-11^ As a RING-type E3 ligase, RNF168 features three main functional domains including a catalytic RING domain as well as two Ub-dependent DSB recruitment modules (UDM1 and UDM2). The N-terminal RING domain engages the nucleosomal H2A–H2B acidic patch and collaborates with the E2 Ub-conjugation enzyme UbcH5c for H2A K13/15 ubiquitylation.^12-14^ The C-terminal UDM2 recognizes H2AK13/15Ub mark for signal propagation or amplification^15^, while the internal UDM1 plays a critical role in the initial recruitment of RNF168 to DSB sites by recognizing K63-linked Ub chains on linker histone H1.^7^ Despite these insights, how RNF168 is modulated by K63-linked ubiquitylated H1 in structurally defined pattern remains unresolved, which is mainly hindered by the difficulty in accessing the homogeneously modified H1 samples.

Recent advances in synthetic protein chemistry have made it an integral part of chromatin research. Methods ranging from total synthesis^16-20^ or semi-synthesis^21-24^ to bioconjugation technology^25, 26^ have been developed to construct a plethora of chemically defined histones with customized post-translational modifications (PTMs), ^16,21,25-30^ which are widely used for many biochemical and biophysical studies to answer mechanistic questions on chromatin regulation. For instance, an engineered sortase variant-mediated chemoenzymatic approach has been developed to generate site-specific acetylated H3 and H2B (e.g., H3K9ac, H3K27ac, H2BK12ac, and H2BK46ac), which enables the biochemical investigations of the activity and selectivity of different histone deacetylases such as HDAC complexes or sirtuin family proteins.^28, 29^ In another example, a semi-synthetic strategy was recently developed to produce phosphorylated and/or ubiquitylated H2AX, which unveiled redundancy in the interplay between histone phosphorylation, ubiquitylation, and methylation on 53BP1 binding.^21^ Despite these advances, the vast majority of contributions made by protein chemistry in the chromatin biochemistry and biophysics area have revolved around core histones (i.e., H2A, H2B, H3, and H4), while technical platforms and chemical tools for linker histone H1 are limited^25, 26, 31^ and need to be further developed to interrogate the role of histone H1 PTMs in epigenetic regulation.

Here we reported the chemical (poly)ubiquitylation of linker histone H1 through a new strategy of CAEPL (Cysteine-Aminoethylation coupled with Enzymatic Protein Ligation), which involves the site-specific alkylation of the recombinant H1 using the bifunctional handle molecule 2-((2-chloroethyl)amino)ethane-1-thiol (CAET),^32, 33^ followed by the Ub-activating E1 enzyme UBA1-assisted protein ligation. Using this strategy, the customized histone H1 with defined ubiquitylated patterns such as mono-ubiquitylation and K63-polyubiquitylation can be readily produced from the recombinant protein precursors in high efficiency and used as tools for the biochemical and structural studies to reveal the *in-trans* crosstalk between H1 ubiquitylation and RNF168-mediated nucleosomal H2A ubiquitylation.

## RESULTS AND DISCUSSION

### Chemical synthesis of ubiquitylated H1.0 with defined ubiquitylation patterns via CAEPL strategy

Previous quantitative proteomics analysis has revealed a significant up-regulation of K63-polyubiquitylation of linker histone H1.0 in response to DSB damage.^6^ Notably, the H1.0 K82 site, located in the globular domain of H1.0 (**Figure 1A**),^34^ exhibited the most pronounced increase in ubiquitylation levels among all the identified H1 ubiquitylated sites.^6^ Therefore, our study commenced with the preparation of the modified H1.0 bearing K63-linked ubiquitin chain at site 82 with different lengths (i.e., H1.0–Ub_n_, n = 1, 2, 3). To achieve these goals, a semi-synthetic route was designed and shown in **Figure 1C**. This strategy can be delineated into three parts: (1) installation of the bifunctional handle 2-((2-chloroethyl)amino)ethane-1-thiol (CAET) at position 82 of H1.0 via cysteine conjugation; (2) enzymatic preparation of K63-linked polyubiquitin chain by the K63-linkage-specific E2 enzyme complex UBC13/MMS2;^35, 36^ and (3) E1-activated native chemical ligation of the CAET-loaded H1.0 with Ub or K63-polyUb.

**Figure 1.**
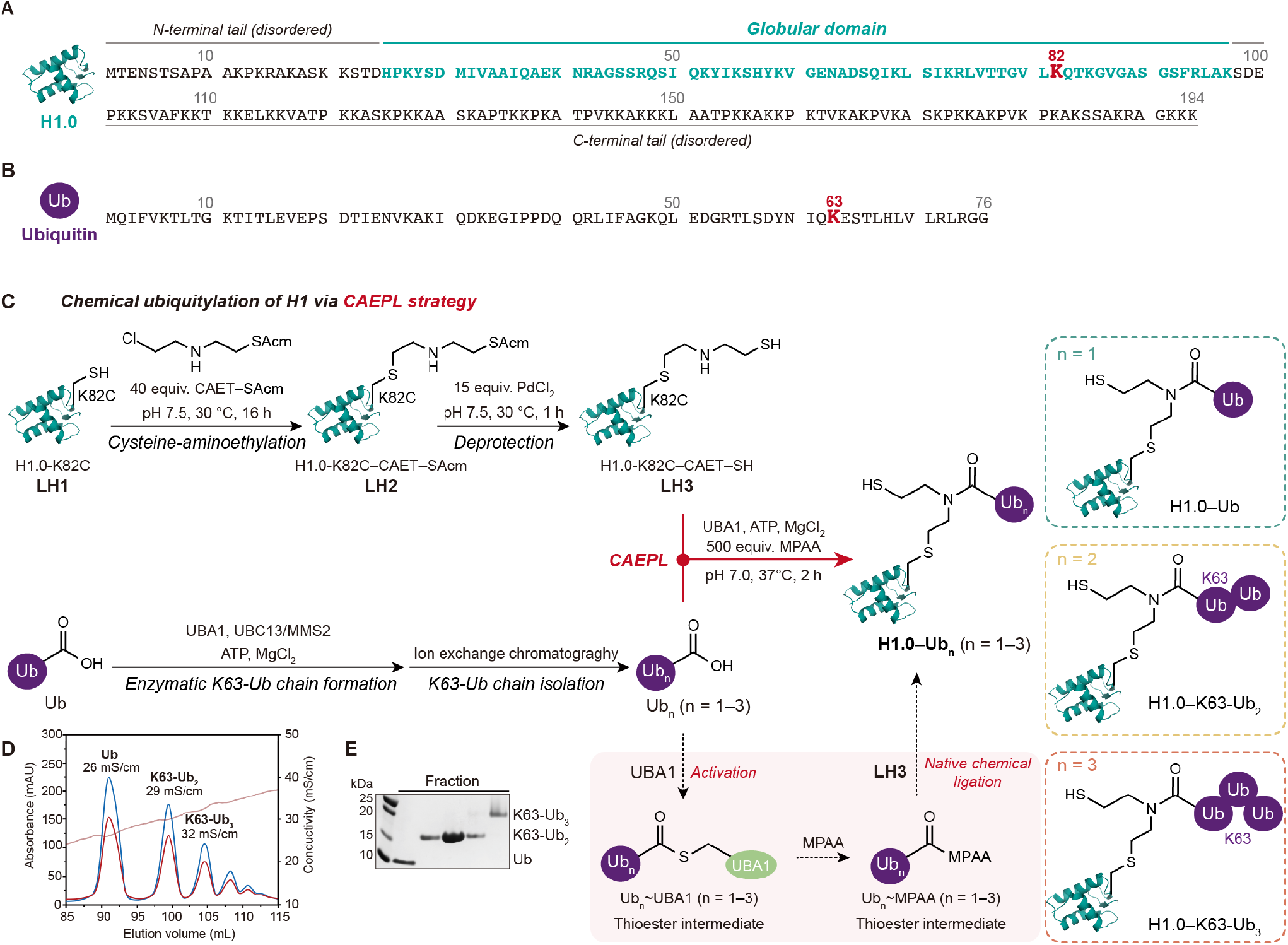
Synthesis of ubiquitylated H1.0 with defined chemical pattern. (**A**) Structural diagram and protein sequence of H1.0. The globular domain of H1.0 was labeled in green and the ubiquitylation site K82 was bolded and colored red. (**B**) Sequence of Ub. The ubiquitylation site K63 for linkage extension was bolded and colored red. The Ub molecule was shown as a purple circle thereafter. (**C**) Schematic route of the Cysteine-Aminoethylation coupled with Enzymatic Protein Ligation (CAEPL) strategy for chemical ubiquitylation of H1.0. Potential intermediates were framed in pink and UBA1 was represent by a light green ellipse. Final products appeared in the dotted box, respectively. (**D**) Cation exchange chromatogram for K63-linked ubiquitin chain mixture generated by UBC13/MMS2-based enzymatic strategy. Blue line indicates 280 nm absorbance of UV detection, red for 260 nm absorbance, and brown for conductivity. (**E**) SDS-PAGE characterization for cation exchange fraction stained with Coomassie brilliant blue.

First, to facilitate the site-specific chemical ubiquitylation of H1.0, the β-mercaptoethylamine group used for native chemical ligation was introduced at K82 by the cysteine-aminoethylation reaction (**Figure 1A**). The H1.0 K82C mutant (**LH1**) was cloned in pET-42b(+) vector and recombinantly expressed in *Escherichia coli* (*E. coli*) with a yield of ca. 20 mg from 1 liter Luria-Bertani (LB) medium. Then, **LH1** was incubated with 2-((2-chloroethyl)amino)ethane-1-(S-acetaminomethyl)thiol (CAET–SAcm, 40.0 equiv.) in unfolding buffer (100 mM PBS, 6 M Gn·HCl, pH 7.5) at 30 °C for 16 hours to give H1.0-K82C–CAET-SAcm (**LH2**). After removal of the S-acetaminomethyl protecting group of **LH2** by treatment with PdCl_2_ (15.0 equiv.) at 30°C for an hour, H1.0-K82C-CAET-SH (**LH3**) was obtained with a yield of 85%.

In parallel, to prepare K63-linked polyubiquitin chains with defined lengths, 1 mM wild-type Ub (the amino acid sequence and K63 site shown in **Figure 1B**) was mixed with 5 μM E2 enzyme complex UBC13/MMS2 in the ubiquitylation buffer (20 mM Tris·HCl, pH 8.0, 150 mM NaCl, 20 mM ATP, 40 mM MgCl_2_), and E1 enzyme UBA1 (1 μM in final concentration) was then added to initiate the enzymatic Ub chain formation. After incubation at 37 °C for 2 hours, the reaction mixture was purified by cation exchange chromatography to give K63-Ub_2_ and K63-Ub_3_, whose identity and purity were characterized by SDS-PAGE analysis (**Figure 1D, E**).

Finally, E1-activated native chemical ligation was performed to assemble H1.0–K63-Ub_n_ (n = 1∼3). **LH3** (200 μM) was mixed with equivalent amounts of Ub, K63-Ub_2,_ or K63-Ub_3_ in the ubiquitylation buffer. Then, an excessive of 4-mercaptophenylacetic acid (MPAA, 500 equiv.) and 2 μM UBA1 were added, followed by pH adjustment to 7.0. The reaction mixture was further incubated at 37 °C for 2–5 hours, and the HPLC monitoring indicated that the condensation of **LH3** with mono-Ub or poly-Ub was observed to be almost quantitative to give the desired H1.0–K63-Ub_n_ (n = 1∼3) products with high efficiency. Notably, in this reaction, activation of Ub or poly-Ub by the E1 enzyme enables the reproducible formation of the thioester intermediates (i.e., Ub_n_∼MPAA, “∼” denotes the thioester bond) for *in situ* native chemical ligation with **LH3**, not only bypassing the additional chemical operations to prepare Ub or poly-Ub thioester but also driving the maximized conversion of protein ligation.

The identity and homogeneity of the protein intermediates (i.e., **LH1, LH2**, and **LH3**) and the desired H1.0-K63-Ub_n_ products (i.e., H1.0–Ub, H1.0–K63-Ub_2,_ and H1.0–K63-Ub_3_) were well characterized by analytical RP-HPLC and electrospray ionization mass spectrometry (ESI-MS) analysis (**Figure 2**). The overall isolated yield was approximately 20% starting from H1.0-K82C material (**LH1**). Using the CAEPL strategy, approximately 3– 5 mg of ubiquitylated H1.0 could be obtained within one week from one liter of *E. coli* recombinantly expressed H1.0-K82C.

**Figure 2.**
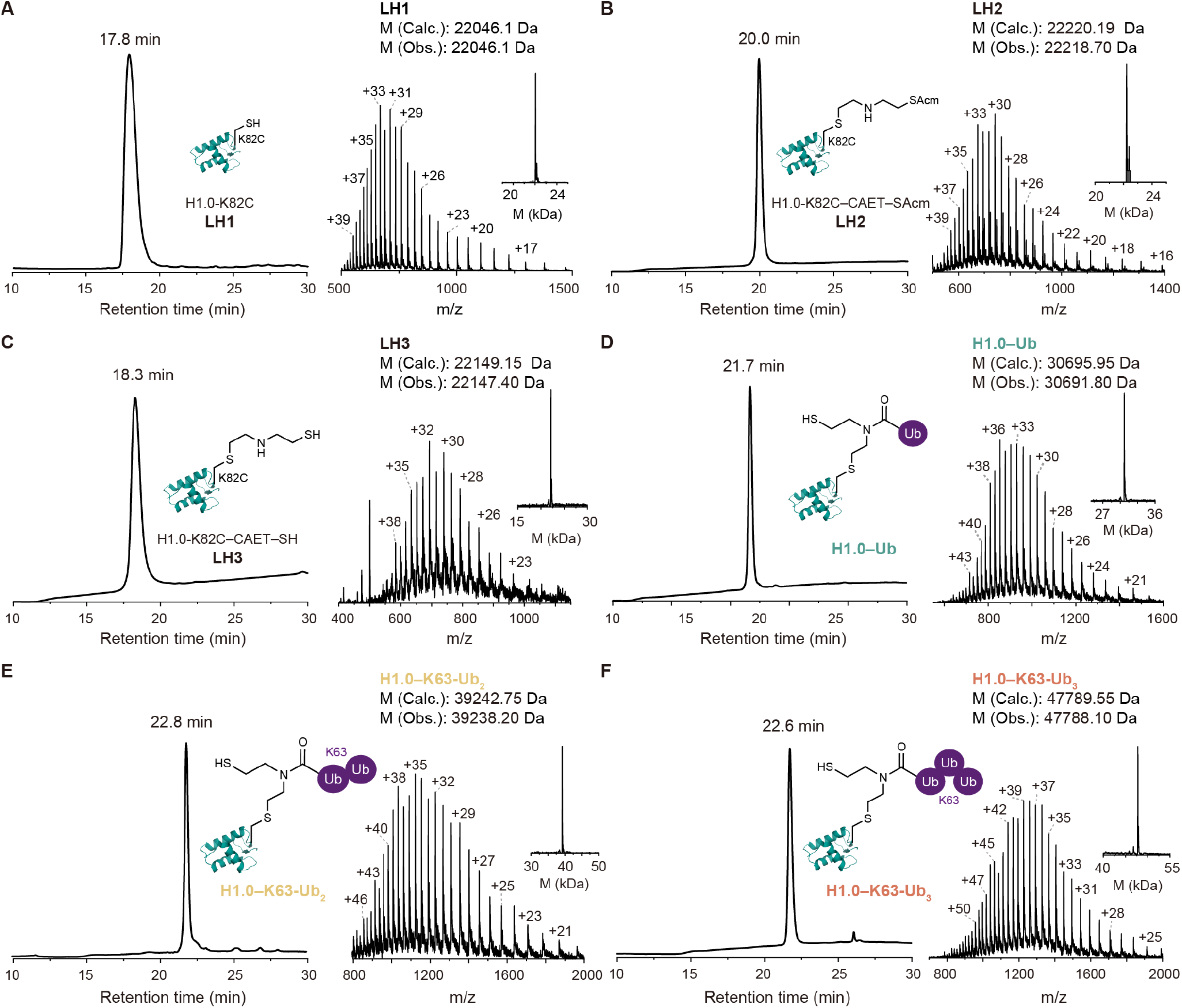
Characterization of the intermediates and the ubiquitylated H1.0 products by HPLC and ESI-MS. (**A**) RP-HPLC (214 nm) and ESI-MS characterization of recombinantly expressed H1.0-K82C (**LH1**). (**B**) RP-HPLC (214 nm) and ESI-MS of H1.0 conjugating CAET–SAcm product H1.0-K82C–CAET–SAcm (**LH2**). (**C**) RP-HPLC (214 nm) and ESI-MS of deprotected H1.0-K82C-CAET-SH (**LH3**). (**D**-**F**) RP-HPLC (214 nm) and ESI-MS characterization of isolated final products H1.0-K82C covalently linked with mono-Ub, K63-linked di-Ub, or tri-Ub, respectively.

### Reconstitution of chromatosomes containing ubiquitylated H1.0

Next, we aimed to incorporate the homogeneously ubiquitylated H1.0 samples into chromatosomes. To this end, four core histones, including the fluorescein-labeled H2A, recombinantly expressed H2B, H3 and H4, were dialyzed against histone refolding buffer (10 mM Tris, 2 M NaCl, 1mM DTT) to give the histone octamers (**Figure S3**). The H2A was labeled with fluorescein at the C-terminus through reaction of H2A-K129C and fluorescein-5-maleimide (CAS: 75350-46-8), which has been previously used for the readily track and detection of RNF168-mediated H2A ubiquitylation assays (see details in **Figure S1**)^37-39^. The histone octamer was then wrapped with 147 bp Widom 601-position DNA with extra 45 bp linker DNA at both the 3’ and 5’ ends (referred to as 45N45 DNA), and dialyzed against the refolding buffer with a gradual decrease of the salt concentration from 2 M to 600 mM. The individual linker histone H1.0 variant (H1.0, H1.0–Ub, H1.0–K63-Ub_2_, or H1.0–K63-Ub_3_) that has been pre-dissolved in the 10 mM HEPES, 600 mM NaCl buffer was added in an equimolar ratio relative to octamer. After 2 hours of static dialysis, the salt concentration of dialysis buffer was gradually reduced from 600 mM to 50 mM by continuous pumping of the HE buffer (10 mM HEPES, 1 mM EDTA). This process resulted in the successful reconstitution of H1.0-containing chromatosomes (**CH0, CH1, CH2**, and **CH3**). We also reconstituted the 45N45 DNA-containing nucleosome core particle (NCP) without linker histone H1.0 (**N**) using the reported method (**Figure 3A**).^40^ All the chromatosomes and NCP were purified by size-exclusion chromatography (SEC) (**Figure 3B**) and characterized by native polyacrylamide gel electrophoresis (Native-PAGE) analysis (**Figure 3C**). Taken together, these results demonstrate the successful assembly of H1.0–K63-Ub_n_ (*n* = 1, 2, 3) chromatosomes (**CH1, CH2**, and **CH3**), unmodified H1.0-containing chromatosome (**CH0**), and 45N45 NCP without linker histone H1.0 (**N**).

**Figure 3.**
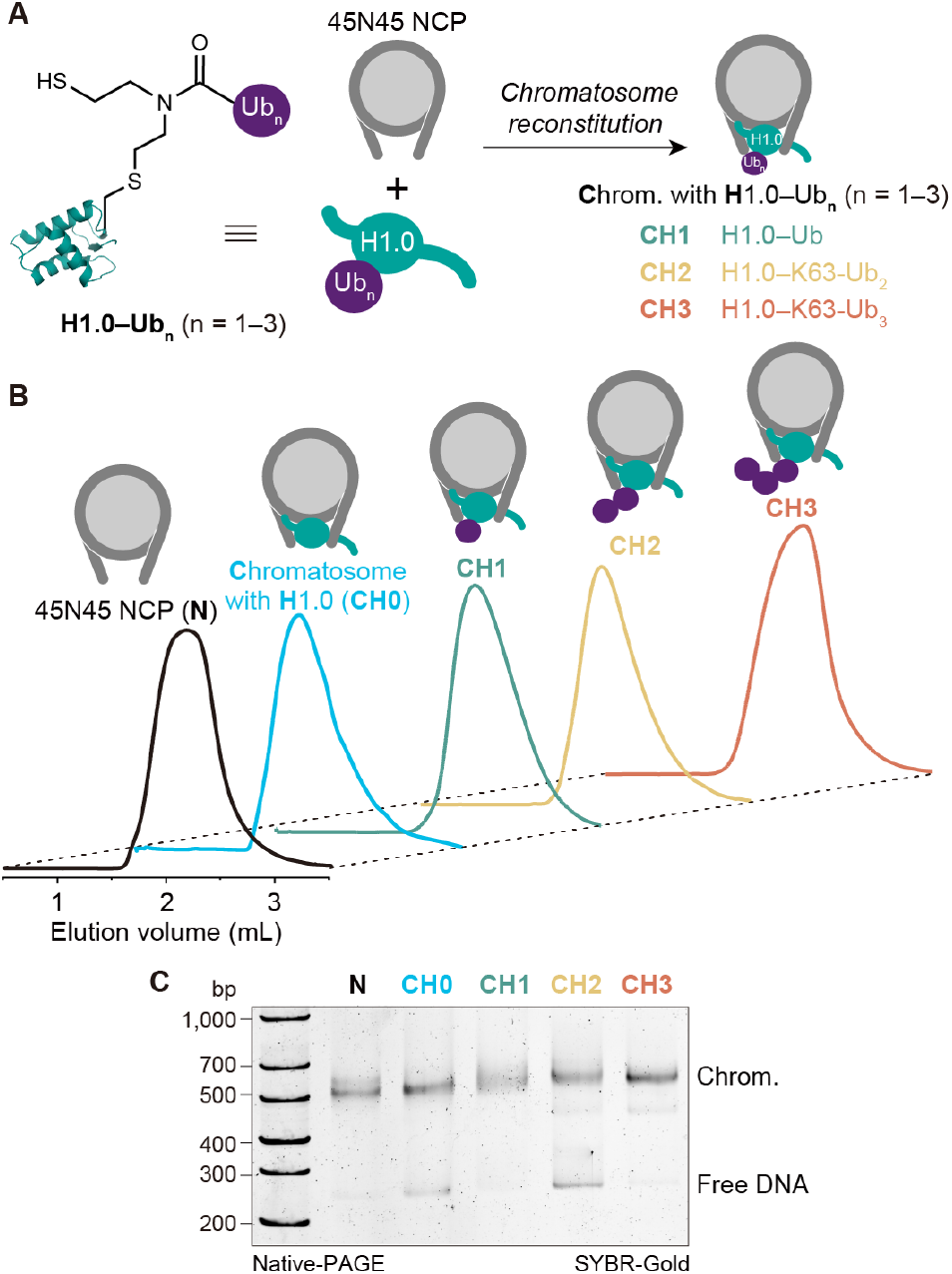
Reconstitution of NCP and chromatosomes containing ubiquitylated H1.0. (**A**) Cartoon representation for chromatosome assembly. (**B**) SEC elution peaks of NCP and chromatosomes. (**C**) Native-PAGE characterization of purified NCP and chromatosomes.

### Stimulation of RNF168-mediated H2A ubiquitylation by K63-linked polyubiquitylated H1.0

With the incorporated nucleosome or chromatosomes in hand, we next set out to understand the role of individual RNF168 structural domains in response to H1.0 ubiquitylation. Four RNF168 constructs (including RNF168^FL^, RNF168^1–193^, RNF168^1–159^, and RNF168^1–113^) were recombinantly expressed and purified (**Figure 4A** and **S4**). *In vitro* RNF168 ubiquitylation assays were conducted using the four RNF168 constructs and tested for activity on nucleosomal or chromosomal substrates of **N, CH0**, and **CH3**. The biochemical results were then analyzed by SDS-PAGE followed by fluorescence imaging and quantification (as shown in **Figure 4B, C**). First, RNF168^FL^ converted about 5% and 10% of H2A to ubiquitylated H2A (H2AUb_n_) on **N** or **CH0** substrates in 5 and 10 min. By comparison, RNF168^FL^ ubiquitylated approximately 40% and 65% of H2A on **CH3** substrate containing H1.0–K63-Ub_3_ in 5 and 10 min, respectively (**Figure 4B, C**), indicating an 8-fold activation of RNF168^FL^ activity by K63-linked tri-Ub on H1.0. Second, similar to RNF168^FL^, the K63-linked tri-ubiquitylated H1.0 can stimulate the ubiquitylation activity of RNF168^1–193^, but not the shorter constructs RNF168^1–159^ or RNF168^1–113^ (**Figure 4B, C**), which suggested that the UDM2 domain is dispensable for polyubiquitylated H1-mediated RNF168 activation, and RNF168^1–193^ containing RING and UDM1 domain was the minimal stimulating construct. Finally, comparing the ubiquitylation results of **N** and **CH0** substrates (**Figure 4B–E**), the presence or absence of unmodified H1.0 barely affects RNF168-mediated H2A ubiquitylation, despite that the RNF168 UDM1 domain can slightly bind unmodified H1.0 in a previous pull-down assay.^7^

**Figure 4.**
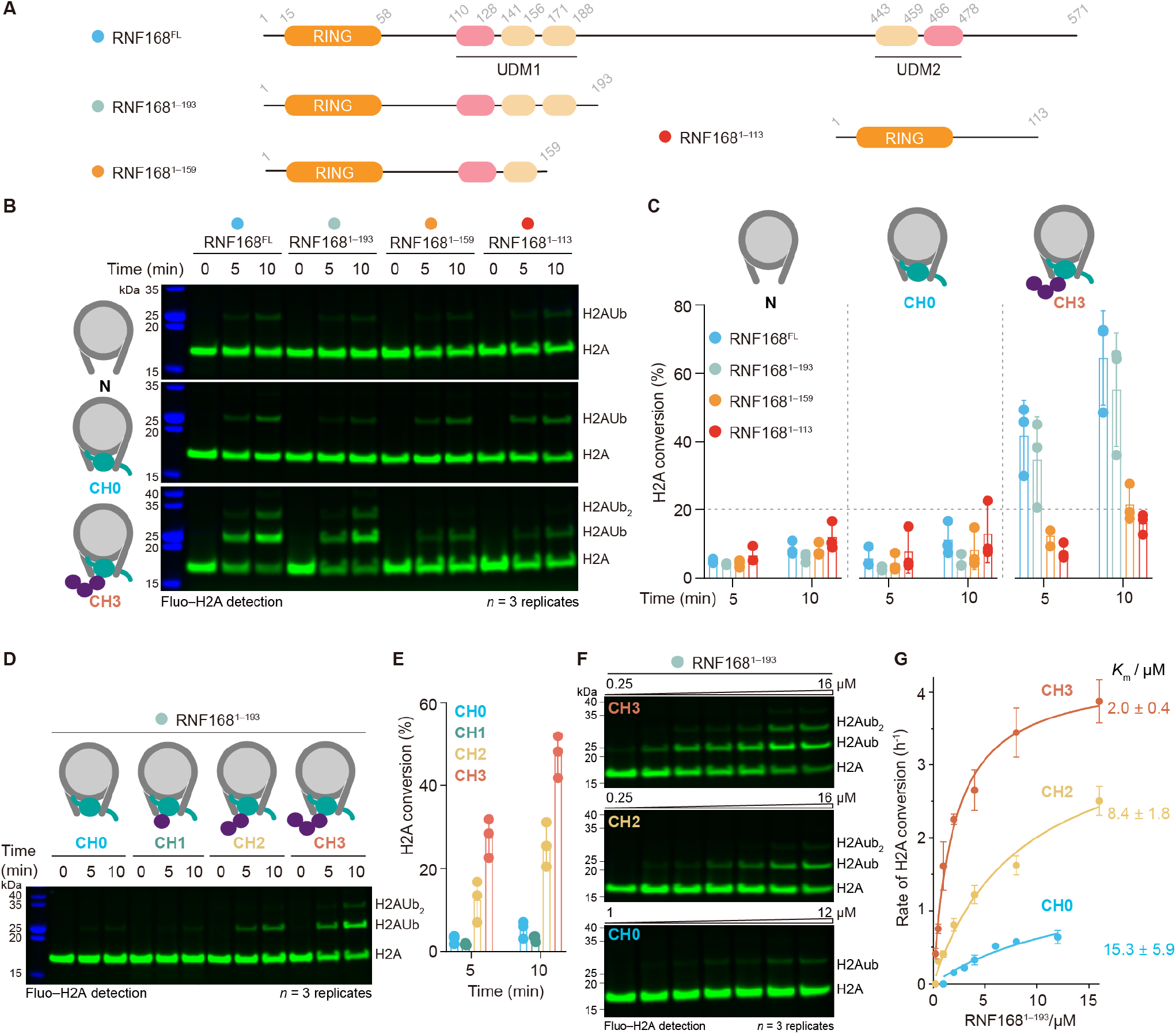
K63-linked polyubiquitylated H1.0 activates RNF168 ubiquitylation activity. (**A**) Domain archetecture of four RNF168 constructs (RNF168^FL^, RNF168^1–193^, RNF168^1–159^, and RNF168^1–113^). The RING domain is responsible for RNF168 ubiquitylation activity. The UDM1 domain contains three motifs related to Ub recognition: LR motif (LRM)1 (residues 110–128), Ub-interacting motif (UMI)1 (residues 141–156), and motif interacting with Ub (MIU)1 (residues 171–188). The UDM2 contains two analogous motifs: MIU2 (residues 443–459) and LRM2 (residues 466–478). (**B**) Fluorescent gels of *in vitro* ubiquitylation assay using the four RNF168 constructs (RNF168^FL^, RNF168^1–193^, RNF168^1–159^, and RNF168^1–113^) on nucleosome (**N**), H1.0 chromatosome (**CH0**) and H1.0–K63-Ub_3_ chromatosome (**CH3**) respectively. (**C**) Quantitative analysis of ubiquitylation activity of the RNF168 constructs on **N, CH0**, and **CH3** in **B**. (**D**) Fluorescent gels of *in vitro* ubiquitylation assay using RNF168^1–193^ on **CH0, CH1, CH2**, and **CH3**, respectively. (**E**) Quantitative analysis of ubiquitylation activity of RNF168^1–193^ construct on **CH0, CH1, CH2**, and **CH3** in **D**. (**F**) Fluorescent gels of RNF168^1–193^ kinetics analysis by titrating RNF168^1-193^ on **CH0, CH2** and **CH3** substrates. (**G**) Three independent experiments were plotted and their averages were fitted to the Michaelis–Menten model to estimate the *K*_m_ constant. Gel images are representative of independent biological replicates (n = 3).

We next investigated how the length of ubiquitin chain on H1.0 modulates RNF168 activity. Four chromatosomes (**CH0, CH1, CH2, CH3**) were used as substrates for RNF168-mediated ubiquitylation assay using RNF168^1–193^, which is activated by H1.0–K63-Ub_3_ to an extent almost comparable to that of RNF168^FL^. As shown in **Figure 4D, E**, gel images read out by fluorescein-labeled H2A showed accelerated H2A ubiquitylation on K63-linked di- or tri-ubiquitylated H1.0 substrates **CH2** (H2A conversion:13% at 5 min; 25% at 10 min) and **CH3** (H2A conversion: 28% at 5 min; 48% at 10 min), but not on monoubiquitylated H1.0-containing **CH1** (H2A conversion: 2% at 5 min; 3% at 10 min), when compared to that of unmodified H1.0-containing **CH0** (H2A conversion: 3% at 5 min; 5% at 10 min). **CH3** exhibited a stronger stimulating activity than that of **CH2** (8-fold versus 5-fold). These data suggested that K63-linked di-Ub on H1.0 is the minimal requirement for RNF168 activation.

To elucidate the enhanced stimulating effect by H1.0–K63-Ub_3_ and H1.0–K63-Ub_2_, we conducted the enzymatic kinetics experiment of RNF168 on chromosomal substrates **CH0, CH2**, and **CH3**. A well-established kinetic method was adopted to determine the apparent Michaelis–Menten constant *K*_m_.^41-43^ Results indicated that the *K*_m_ values of RNF168^1–193^ on **CH0, CH2**, or **CH3** were 15.3 ± 5.9 μM, 8.4 ± 1.8 μM, and 2.0 ± 0.4 μM, respectively (**Figure 4G**), suggesting an enhanced binding affinity of RNF168^1-193^ for **CH2** or **CH3** compared to **CH0**. Notably, the decrease in *K*_m_ value of **CH2** and **CH3** correlates with the increment in the number of K63-linked ubiquitin molecules on H1.0. The RNF168 UDM1 domain has been reported to bind K63-linked di-Ub. Thus, the longer K63-linked tri-Ub provides an additional binding opportunity for UDM1, which may explain the higher affinity and activity of RNF168 for **CH3** substrate.

### Structure analysis of RNF168/Ubch5c module bound to K63-ubiquitylated H1.0 chromatosome

To investigate the mechanism of how RNF168 recognizes H1.0–K63-Ub_n_ on the chromatosomes to stimulate its H2A ubiquitylation activity, we set out to analyze the structure of RNF168/UbcH5c-bound H1.0–K63-Ub_3_ chromatosomes complex. A previously developed chemical trapping strategy was utilized to obtain the complex.^12, 37, 44^ In detail, we synthesized an H2A ubiquitylation mimic **HA1**, which covalently attaches the H2A K15 site and the Ub carboxyl terminus to the UbcH5c active Cys residue via a CAET handle (**Figure 5A**). Next, **HA1** was mixed with three core histones H2B, H3, and H4 to reconstitute a histone octamer, which was then assembled with 45N45 DNA and H1.0–K63-Ub_3_ to form the chromatosome probe **CU3** (**Figure 5A**). Denatured SDS-PAGE analysis showed that both the H2A ubiquitylation mimic **HA1** and H1.0–K63-Ub_3_ were incorporated into the chromatosome probe **CU3** (**Figure 5A**). We have demonstrated that the H2A ubiquitylation stimulation effect by H1.0–K63-Ub_3_ of the RNF168^1–193^ fragment containing the RING and the UDM1 domain was comparable to that of RNF168^FL^ (**Figure 2B, C**). Therefore, we incubated RNF168^1–193^ with the chromatosome probe **CU3** to assemble the **CUP3** complex (**Figure 5A**). The **CUP3** complex was glutaraldehyde cross-linked, purified by SEC, and subjected to single-particle cryo-EM analysis. A cryo-EM reconstruction at an overall resolution of 4.3 Å was obtained (**Figure 5B, C, Figure S5**, and **Table S1**). In this **CUP3** cryo-EM reconstruction, we were able to assign the densities of the core histone octamer, ∼170 bp DNA, and the linker histone H1.0 (**Figure 5B, C**, and **S6**). The **CUP3** cryo-EM reconstruction matches well with a reported structure of H1.0-containing chromatosome (PDB 5NL0).^34^ The H1.0 globular domain extensively engages the SHL0 DNA and the linker DNA of the chromosome (**Figure 5B** and **S6**), consistent with the well-established model that H1 locates at the chromosomal dyad.^45, 46^ The RNF168 RING domain associates with the nucleosomal H2A–H2B acidic patch through an α-helix (residues 63– 68), orientating UbcH5c towards the H2A K13/15 (**Figure B, C**). The Ub motif of H2A K15 is located between the back of UbcH5c and DNA SHL 3.5, consistent with the reported structures of the RNF168/Ubch5c–Ub/NCP complex (**Figure 5B, C**),^12, 14^ suggesting that the RNF168 RING/UbcH5c ubiquitylation module binds to chromatosomes containing K63-linked tri-ubiquitylated H1.0 in almost the same manner as it binds to nucleosomes without H1.0.

**Figure 5.**
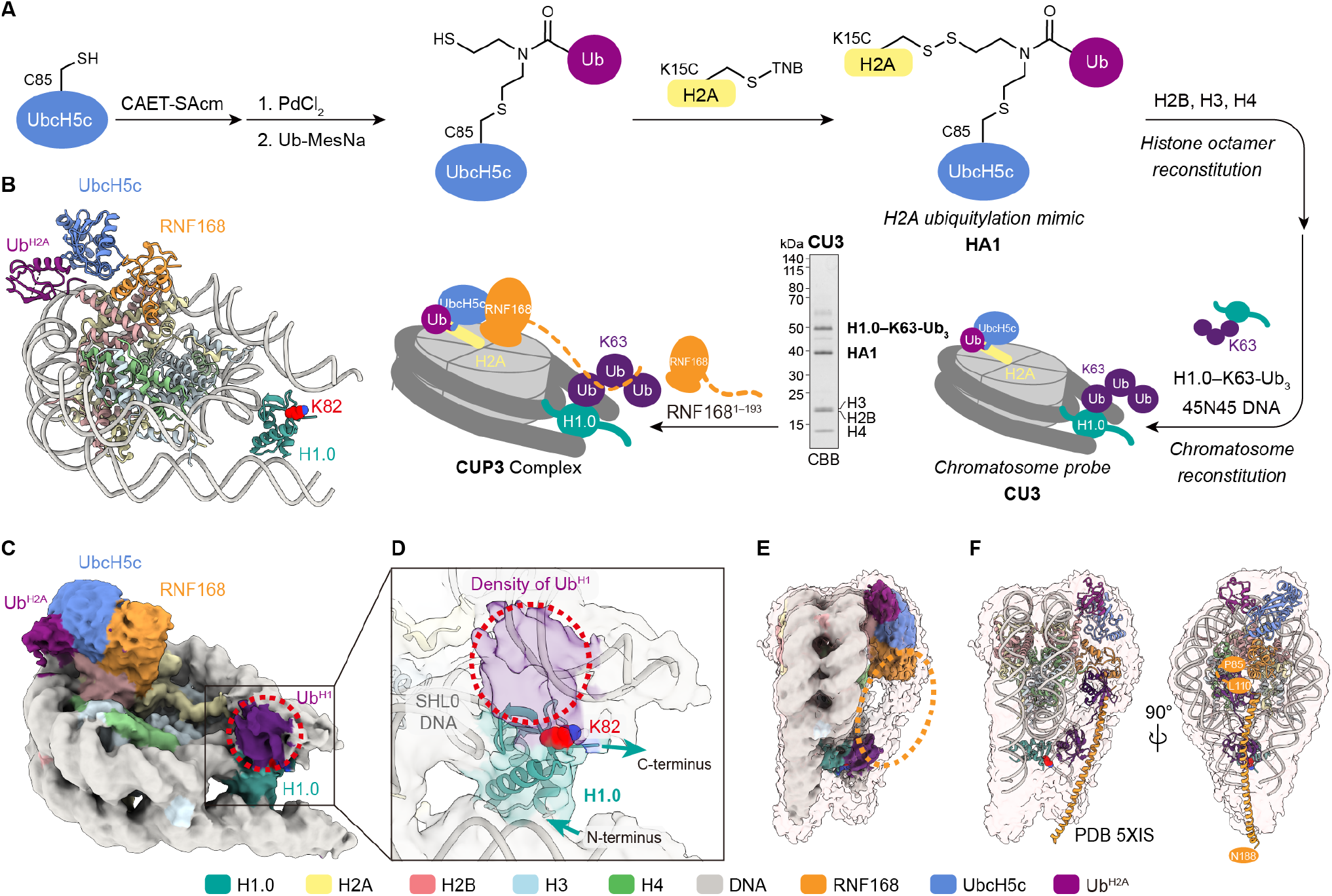
Structure of RNF168/Ubch5c module bound to H1.0–K63-Ub_3_ chromatosome. (**A**) Synthetic route of the H2A ubiquitylation mimic HA1 and workflow for preparing the RNF168^1–193^/Ubch5c– Ub/ H1.0–K63-Ub_3_ chromatosome (**CUP3**) complex. (**B**) Model of **CUP3** complex. (**C**) Cryo-EM density map of **CUP3** complex at the threshold level of 0.0064. (**D**) Close-up view of the cryo-EM density of H1.0 and the extra density for Ub^H1^ above H1.0 K82 site. (**E**) Cryo-EM density map of **CUP3** complex in a semi-transparent molecular envelope (threshold level of 0.0030). A stretch of density between RNF168 RING and H1.0 is circled by an orange dashed oval. (**F**) Different views of the **CUP3** complex model and RNF168 UDM1–K63-Ub_2_ crystal structure (PDB 5XIS) fitted in a semi-transparent molecular envelope (threshold level of 0.0030). The RNF168 RING C-terminus (residue P89), UDM1 N-terminus (residue L110), and UDM1 C-terminus (residue N188) are indicated.

Notably, nucleosomes have pseudo-C2 symmetry^47^ and binding of linker histone H1.0 disrupts this symmetry, resulting in a difference in the disc surfaces on either side of the nucleosome (**Figure 5C, D**). Interestingly, in the **CUP3** complex structure, the H1.0 K82 site is on the same side of the nucleosome disc surface where the RNF168/UbcH5c ubiquitylation module binds, suggesting that K63-Ub_3_ conjugated to H1.0 K82 site recruits RNF168 to the same side of the nucleosome disc surface (**Figure 5C, E**). There is an extra density above the K82 site of H1.0, which might be attributed to the density of Ub^H1^ motif conjugated to H1.0 K82 (**Figure 5C, D**). When the density map threshold level is lowered from 0.0064 to 0.0030, a continuous stretch of density is observed between RNF168 and H1.0 (**Figure 5E**). We speculate that this stretch of density may correspond to the recognition of H1.0–K63-Ub_3_ by RNF168 residues (especially UDM1). A previously reported crystal structure of RNF168 UDM1 helix in complex with K63-linked diUb (PDB: 5XIS),^48^ is well accommodated by this stretch of density (**Figure 5F**). In this docking model, the RNF168 UDM1 N-terminus (residue L110) is orientated towards the RING C-terminus (residue P89), which are about 15 Å apart (**Figure 5F**). The length of an amino acid between RING and UDM1 (residues 90–109) can satisfy this distance.

### A comparable RNF168 stimulation effect of poly-ubiquitylated H1.0 in both the inter- and intra- nucleosomal crosstalk

We have shown that K63-linked polyubiquitylated H1.0 can effectively stimulate the H2AK13/15 ubiquitylation of mono-nucleosomes by RNF168 (**Figure 4**). It should be noted that the H1 ubiquitylation couples initiation and amplification of the H2AK13/15Ub signal in response to DNA damage *in vivo*.^6, 7^ This poses a question: can the polyubiquitylated H1 augment the RNF168-mediated ubiquitylation in an inter-nucleosomal manner? To investigate this possibility, we carried out the RNF168 ubiquitylation assays on an asymmetric H1.0- polyubiquitylated di-nucleosome substrate (**Figure 6**).

**Figure 6.**
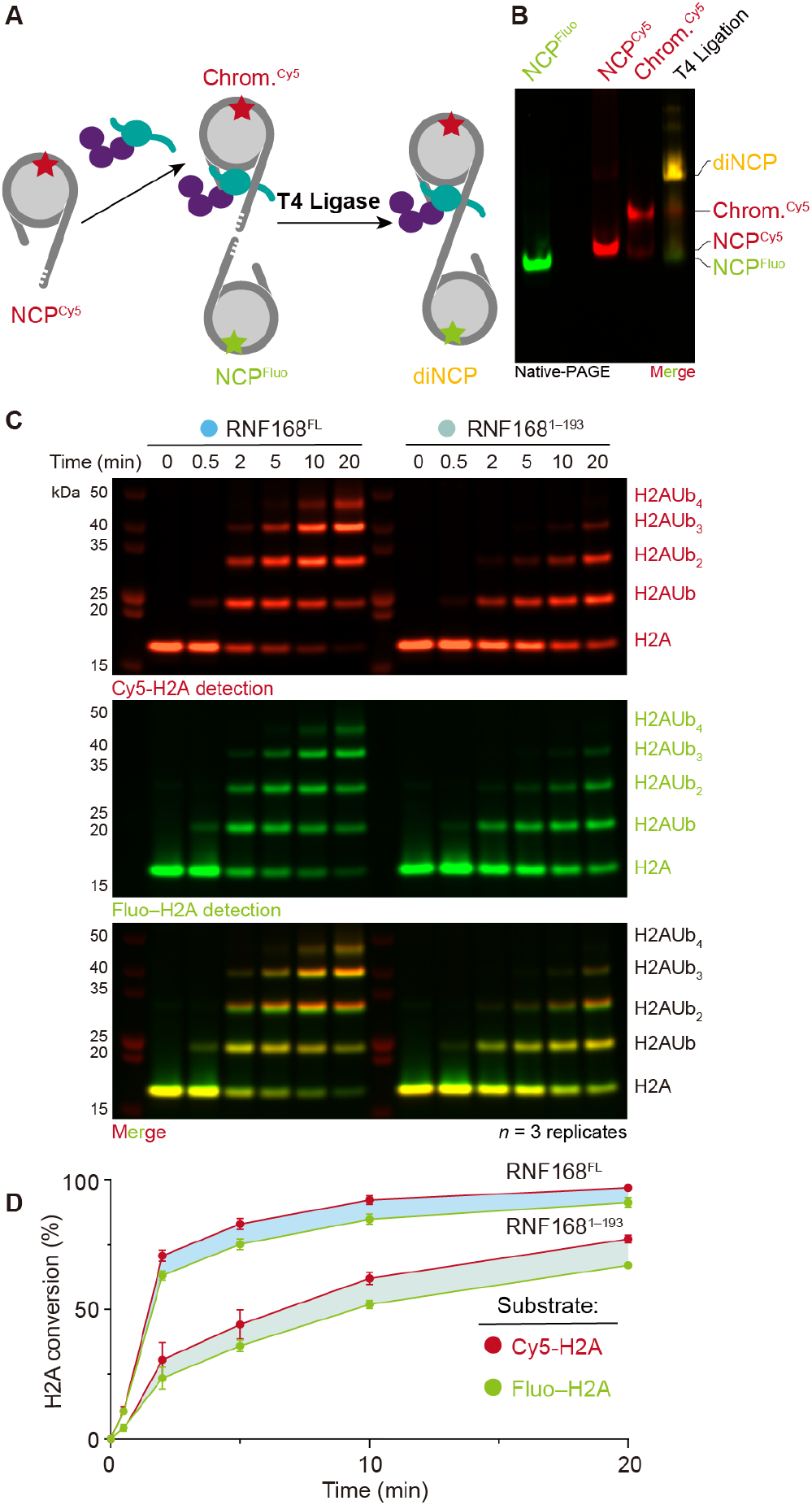
Reconstitution of asymmetric di-nucleosome and ubiquitylation assay with RNF168. (**A**) Schematic depiction of asymmetric di-nucleosome reconstitution. Mono-nucleosome Chrom^Cy5^ and NCP^Fluo^ were assembled with different sticky end DNA and fluorescent molecular covalently conjugated octamer. T4 ligase was used for mono-nucleosome ligation generating asymmetric di-nucleosome. (**B**) Characterization of mononucleosomes and T4 ligation for di-nucleosome by 4.5% native-PAGE, visualized in red or green channel and two channels merged one. (**C**) RNF168^FL^ (left) and RNF168^1-193^ (right) ubiquitylation assay on di-nucleosome analyzed by SDS-PAGE and detected by fluorescence imaging. (**D**) Quantitative analysis of di-NCP ubiquitylation assay. Red plot represents Chrom^Cy5^ and green one for NCP^Fluo^. Horizontal differentiation between different mono-NCP for RNF168^FL^ and RNF168^1-193^ was labeled with light sky blue and dark sea green, respectively.

The core histone H2As on each NCP unit of the di-nucleosome were labeled with Cy5 (Cy5–H2A, detected in red channel) or fluorescein (Fluo–H2A, detected in green channel) respectively for a separated visualization in ubiquitylation assays (the synthetic characterization of labeling was shown in **Figure S1, S2**). To reconstitute the asymmetric di-nucleosome, we first reconstituted the Cy5–H2A chromatosome containing H1-K63-Ub_3_ (Chrom.^Cy5^), and Fluo–H2A nucleosome (NCP^Fluo^). It should be noted that two Widom 601 DNAs with complementary overhangs at the DNA ends were used for the indicated assembly (see **Supplementary Methods** for details). Then the T4 ligase-mediated ligation of the Chrom.^Cy5^ and NCP^Fluo^ was conducted based on the complementary paired sticky-end DNAs to give the asymmetric H1-K63-Ub_3_ modified di-nucleosome (**Figure 6A**). The reconstituted Chrom.^Cy5^ and NCP^Fluo^ were characterized by native-PAGE visualized in red or green channels, and a discrete band that simultaneously emitted fluorescence at binary channels migrated over both Chrom.^Cy5^ and NCP^Fluo^ bands in native-PAGE, indicating the formation of a ligated di-nucleosome (**Figure 6B**).

The asymmetric di-nucleosome was then subjected to RNF168 ubiquitylation assays using RNF168^FL^ and RNF168^1–193^ constructs and results were visualized in red or green channels. RNF168^FL^ showed a significant higher ubiquitylation activity than RNF168^1-193^, and remarkably, both RNF168^FL^ and RNF168^1–193^ exhibit comparable efficiency in ubiquitylating the Cy5–H2A of the Chrom.^Cy5^ containing H1-K63-Ub_3_ in comparison to the Fluo–H2A of the NCP^Fluo^ (**Figure 6 C, D**). It is speculated that the K63-linked tri-ubiquitylated H1 may recruit the RNF168 UDM1 domain, while exhibiting a relatively high degree of structural flexibility on the chromatosomes, and can target the N-terminal catalytic RING domain to ubiquitylate the intra-nucleosomal or inter-nucleosomal H2A K13/15 sites.

## CONCLUSION

In summary, we have developed an expedient and convergent strategy termed CAEPL, which uses the bifunctional handle molecule CAET to integrates the cysteine aminoethylation reaction and the E1-activated native chemical ligation to prepare homogeneously modified H1.0 with defined ubiquitylation patterns. Using this strategy, we successfully obtained monoubiquitylated and K63–linked di- and tri-ubiquitylated H1.0 histone, which could be incorporated into individual chromatosomes. Quantitative biochemical analysis of different RNF168 constructs on substrates including NCP, H1.0 chromatosome, H1.0–Ub chromatosome, H1.0–K63-Ub_2_chromatosome and H1.0–K63-Ub_3_ chromatosome demonstrated that K63–linked poly-ubiquitylated H1.0 could directly stimulate RNF168 ubiquitylation activity by enhancing the affinity of RNF168 for chromatosome. Subsequent cryo-EM structural analysis of the RNF168/UbcH5c–Ub/H1.0–K63-Ub_3_ chromatosome complex revealed the potential recognition of RNF168 UDM1 domain for K63-linked polyubiquitylation on H1. We also investigated the biochemical behavior of RNF168 on asymmetric di-nucleosome involving intra- and inter-nucleosomal activity. Collectively, our CAEPL strategy provides chemically-defined (poly)ubiquitylated linker histone H1.0 to study the crosstalk between H1 ubiquitylation and H2A ubiquitylation mediated by RNF168, shedding new light on a more comprehensive understanding of the PTM regulatory network. It is expected that the CAEPL strategy as a general approach for preparing homogeneously (poly)ubiquitylated proteins may find more applications to the biochemical and biophysical studies on the ubiquitylation of other protein families.

## SUPPORTING INFORMATION

General reagents and experimental procedures; Cloning and plasmid construction; Protein expression and purification; Chemical ubiquitylation of H1.0 via the Cysteine-Aminoethylation coupled with Enzymatic Protein Ligation (CAEPL); Preparation of Fluorescein-labeled H2A and Cy5-labeled H2A; DNA preparation; Reconstitution of histone octamers, nucleosomes, chromatosomes and asymmetric di-nucleosome; *In vitro* ubiquitylation assays; Cryo-EM sample preparation; Cryo-EM data collection and processing; Model building.

## NOTES

The authors declare no competing financial interest.

## ACKNOWLEDGMENT

We thank the National Key R&D Program of China (grant nos. 2022YFC3401500) for financial support. This study was supported by the National Natural Science Foundation of China (grant nos. 22137005, 92253302, 22227810). This work has been supported by the XPLORER prize and the New Cornerstone Science Foundation. We acknowledge the Tsinghua University Branch of China National Center for Protein Sciences Beijing for cryo-EM data collection.

