## Supplementary Information for "Promotion of RNF168-Mediated Nucleosomal H2A Ubiquitylation by Structurally-defined K63-Polyubiquitylated Linker Histone H1"

### Table of Contents

**1. General……………………………………………………………………………..3**

**1.1 General Reagents ……………………………………………………………..3**

**1.2 HPLC, SDS-PAGE, Native PAGE and Mass Spectrometer……………………..…3**

**2. Cloning and plasmid construction……………………………………………………..4**

**3. Protein expression and purification……………………………………………………4**

**4. Chemical ubiquitylation of H1.0 via the Cysteine-Aminoethylation coupled with Enzymatic Protein Ligation (CAEPL)……………...…………………..…………………………………...4**

**4.1 Preparation of H1.0–K82C–CAET–SH (LH3)……………..……………………4**

**4.2 Preparation of K63-linked Ubiquitin Chains……………………………………5**

**4.3 Preparation of H1.0–K63-Ub$_n$ (n = 1–3) via the Cysteine-Aminoethylation coupled with Enzymatic Protein Ligation (CAEPL)…………………..………………………5**

**5. Preparation of Fluorescein-labeled H2A and Cy5-labeled H2A……………………………5**

**6. DNA Preparation…………………………………………………..……………………..5**

**7. Reconstitution of histone octamers, nucleosomes, chromatosomes and asymmetric di-nucleosome……………………………………………………………………………..........…6**

**8. *In vitro* ubiquitylation assays………………………………………………….…...……..7**

**8.1 Ubiquitylation assays on nucleosome, H1.0 chromatosome and H1.0–K63-Ub$_n$ chromatosomes…………….....………………………………………….…………………7**

**8.2 Kinetics assays……………………….…………………………………………8**

**8.2 Ubiquitylation assays on asymmetric di-nucleosome……….…………………………8**

**9. Cryo-EM sample preparation……………………………………………………………..8**

**10. Cryo-EM data collection and processing…………………………...………...…9**

**11. Model building……………………………………………………………………...…..9**

**Supplementary References………………………………………………………….…10**

**Figure S1–6………………………...…………………………………………………...…11**

**Table S1……………………………………………………………………………………15**

### 1. General

#### *1.1 General Reagents*

Guanidine hydrochloride (Gn · HCl), ethylenediaminetetraacetic acid (EDTA), tris(hydroxymethyl) aminomethane (Tris), sodium chloride (NaCl), 2-[4-(2-hydroxyethyl)piperazin-1-yl]ethanesulfonic acid (HEPES) and tris(2-carboxyethyl)phosphine hydrochloride (TCEP·HCl) were purchased from Adamas-beta (Shanghai, China). Acetonitrile (HPLC grade) and trifluoroacetic acid (TFA, HPLC grade) were purchased from J. T. Baker (Phillipsburg, NJ, USA). 4-mercaptophenylacetic acid (MPAA) was purchased from Alfa Aesar. Cy5 maleimide and Fluorescein-5-maleimide were purchased from Aladdin (Shanghai, China). Chemically competent Trans5α and BL21 (DE3) cells were purchased from TransGene Biotech (Beijing, China). Glutathione-sepharose resin and Ni-NTA resin were purchased from LABLEAD (Beijing, China). Superdex75/200 10/30 GL column, Superose 6 Increase 5/150 GL column, and Source 15S/15Q 10/30 GL column were purchased from GE Healthcare. Codon optimization and gene synthesis services were purchased by GenScript Biotech (Nanjing, China). The T4 ligase was purchased from LABLEAD (Beijing, China).

#### *1.3 HPLC, SDS-PAGE, Native PAGE, FPLC, and Mass Spectrometer*

Reversed-phase HPLC (RP-HPLC) was performed on Shimadzu Prominence HPLC. Analytical Welch XB-C4 (4.6 × 250 mm, 5 μm particle size) or C18 (4.6 × 250 mm, 5 μm particle size) columns were used at a flow rate of 1.0 mL/min for analysis. Semi-preparative Welch XB-C4 (21.2 × 150 mm, 5 μm particle size) and C18 (10× 250 mm, 5 μm particle size) columns were used at a flow rate of 4 mL/min for peptide semi-preparation. Buffers for RP-HPLC: buffer A (0.1% TFA in $CH_3CN$) and buffer B (0.1% TFA in water). Both solvents were sonicated for 20 min before use. The UV absorption was monitored at 214nm and 254 nm.

For SDS-PAGE, samples were loaded onto 12% NuPAGE Bis-Tris gels (Thermo Fisher Scientific) or 4-12% SurePAGE Bis-Tris gels (GenScript) and electrophoresed for 25min at 160 V. Pictures were taken on the ChemiDoc$^{TM}$ XRS+ system (Bio-Rad).

For Native PAGE, samples were loaded onto 4.5% Native PAGE gels and electrophoresed at 180 V for 45min at 4°C. Gels were stained with SYBR-Gold dye (Thermo Fisher Scientific) for 10min and visualized on the ChemiDoc$^{TM}$ XRS+ system (Bio-Rad).

FPLC was performed on an ÄKTA Purifier (GE Healthcare Life Science). All the buffers were filtered through 0.22 μm filter paper. Every injection was monitored at the wavelength of 280 nm, 260 nm, and 214 nm.

ESI-MS was measured on LC/MS 2020 (SHIMADZU).

### 2. Cloning and plasmid construction

RNF168$^{FL}$, RNF168 fragments (RNF168$^{1–113}$, RNF168$^{1–159}$, RNF168$^{1–193}$), and UbcH5c were cloned into plasmids as previously described.[1] Human UBA1, Ub, and individual human core histones (including H2A, H2B, H3(C96S/C110S), and H4) were cloned as previously described.[2] The cDNA of human linker histone H1.0 K82C was synthesized and sequence was optimized by GenScript Biotech (Nanjing, China) for Escherichia coli overexpression. The H1.0 gene (Human, uniport ID: P07305) was cloned into the pET-42b(+) vector with a C-terminal 8xHis tag. Truncations and mutations were generated by homologous recombination or standard site-directed PCR mutagenesis methods.

### 3. Protein expression and purification

RNF168$^{FL}$, RNF168 fragments (RNF168$^{1–113}$, RNF168$^{1–159}$, RNF168$^{1–193}$), UbcH5c, Ub, UBA1, and individual human core histones were expressed in BL21(DE3) *Escherichia coli* (*E. coli*) cells and purified as previously described.[1] For H1.0-K82C, monoclonal colonies containing the desired H1.0-K82C gene were screened on kanamycin plates. Single colonies from the plates were transferred to a 10 mL Luria-Bertani (LB) medium containing 50 μg/mL kanamycin and grown overnight at 37°C. The pre-culture was added to 1 L LB medium (supplemented with 50 μg/mL kanamycin) and further grown at 37°C until an $OD_{600}$ of 0.6-0.8. Induction of H1.0 K82C protein expression in *E. coli* was achieved by adding 1 mM IPTG, followed by incubation at 37°C for 3 h and centrifugation at 4,000 rpm to collect cell pellets. Then the cell pellets were resuspended in Lysis Buffer (20 mM HEPES, 750 mM NaCl, 8 mM DTT, 6M Gn·HCl, pH 7.8) and subjected to sonication at 120 W for 30 minutes. The lysate was ultracentrifugation at 12,000 rpm for 30 min at 4°C and the supernatant was loaded onto a Ni-NTA affinity resin. H1.0-K82C was eluted by H1 Lysis Buffer supplemented with 400 mM imidazole and further purified by reversed-phase high-performance liquid chromatography (RP-HPLC) and lyophilized into powder.

### 4. Chemical ubiquitylation of H1.0 via the Cysteine-Aminoethylation coupled with Enzymatic Protein Ligation (CAEPL)

#### *4.1 Preparation of H1.0–K82C–CAET–SH (LH3)*

H1.0-K82C (**LH1**) powder was dissolved in unfolding buffer (100 mM PBS, 6 M Gn·HCl, pH 7.5), followed by the addition of 40 equivalents of CAET–SAcm molecule (1 M stock solution in DMSO). The pH was adjusted to 7.5, and the mixture was reacted at 30°C for 16 h. After semi-preparative HPLC purification, the reaction product H1.0–K82C–CAET–SAcm (**LH2**) was obtained and lyophilized into powder. **LH2** powder was then dissolved in unfolding buffer to a final concentration of 1 mM. Subsequently, 15 equivalents of $PdCl_2$ (100 mM in 6M Gn·HCl) and 5mg/mL TCEP were added, and the pH was adjusted to 7.5. After reacting at 30°C for an hour, analysis by HPLC-MS confirmed complete removal of the S-acetamidomethyl (Acm) protecting group (converted entirely to the product with a molecular weight decrease of 71). Semi-preparative HPLC purification combined with mass spectrometry confirmed the molecular weight identification of H1.0–K82C–CAET–SH (**LH3)**, which was finally lyophilized into powder.

#### *4.2 Preparation of K63-linked Ubiquitin Chains*

For K63-linked ubiquitin chain generation, 1 μM hUBA1, 5 μM Ubc13/Mms2 and 1 mM Ub were combined in the ubiquitylation buffer (20 mM Tris·HCl, pH 8.0, 150 mM NaCl, 20 mM ATP, 40 mM $MgCl_2$), and the reaction was carried out at 37°C for 2 hours. The reaction was quenched with 5 volumes of buffer A (50 mM NaOAc, pH 4.5), followed by ion exchange purification and identification by SDS-PAGE to collect K63-$Ub_2$ (29.9 mS/cm) and K63-$Ub_3$ (32.4 mS/cm) fractions, which were subsequently dialyzed into 20 mM HEPES, 150 mM NaCl, pH 7.5.

#### *4.3 Preparation of H1.0–K63-Ub$_n$ (n = 1–3) via the Cysteine-Aminoethylation coupled with Enzymatic Protein Ligation (CAEPL)*

In the ubiquitylation buffer (20 mM Tris·HCl, pH 8.0, 150 mM NaCl, 20 mM ATP, 40 mM $MgCl_2$), 200 μM H1.0–K82C–CAST–SH (**LH3)** powder was dissolved in the buffer (20 mM Tris, pH 7.5, 150 mM NaCl) with the equivalent Ub, K63-linked $Ub_2$ or $Ub_3$. Additionally, 100 mM MPAA was supplemented, first dissolving MPAA powder at pH 5, then adjusting to pH 7.0, then adding UBA1 to a final concentration of 2 μM. The reaction was carried out at 37°C for 2 hours. After semi-preparative HPLC purification and mass spectrometry identification, pure H1.0–K63-$Ub_n$ (n = 1~3) sample was obtained and lyophilized. Deconvolution for molecular weight was calculated by GUI_UniDec.[3]

#### 5. Preparation of Fluorescein-labeled H2A and Cy5-labeled H2A

1 μM H2A-K129C peptide was dissolved in unfolding buffer (100 mM PBS, 6 M Gn·HCl, pH 7.5) and fluorescein-5-maleimide or Cyanine5 maleimide (1.2 μM) was added into buffer. The reaction was carried out at 37°C for 2 hours and then purified by semi-preparative HPLC followed by ESI-MS characterization.

#### 6. DNA Preparation

The 237-bp 45N45 DNA for nucleosome or chromatosome was prepared by PCR amplification using a synthesized 45N45 DNA (single copy) pUC19 plasmid and purified by ion exchange. Sequences of 45N45 DNA is listed below (the sequence of Widom 601 DNA is underlined):

ATCAAAGCTGGGTACCAGTTCTGAGATCACCCTAGGTCTCTGATGCTGGAGAATCCCGGTGCCGAGGCCGCTCAATTGGTCGTAGACAGCTCTAGCACCGCTTAAACGCACGTACGCGCTGTCCCCCGCGTTTTAACCGCCAAGGGGATTACTCCCTAGTCTCCAGGCACGTGTCAGATATATACATCCTGTCACGCGGTGAACAGCGAGATCGGATACACTAGTTCTAGAGCGGAT

To obtain DNA 21N16 and 16N3 with sticky ends, different lengths of DNA linker at both ends of the 601 sequence DNA, including DNA 21N33 and 30N3 with a DraIII restriction enzyme site on one side were amplified by PCR using 30N30-pUC19 plasmid as a template. The obtained DNA was purified using anion exchange mono Q (A Buffer: 10 mM HEPES, 1 mM EDTA, pH 7.5; B Buffer: 10 mM HEPES, 1M NaCl, 1 mM EDTA, pH 7.5) and dialyzed overnight into EDTA free buffer (10 mM HEPES, pH 7.5). Each DNA was treated with DraIII (NEB, R3510L) at a ratio of 1 μg DNA to 5 units DraIII, followed by the addition of 1/10 volume of 10x rCutSmart Buffer (500 mM Potassium Acetate; 200 mM Tris-acetate; 100 mM Magnesium Acetate; 1 mg/ml Recombinant Albumin; pH 7.9), and incubated at 37°C for 1 hour. The DNA was then purified using a mono Q column to remove DraIII, resulting in DNA 21N16 and 16N3 with sticky ends. Sequences of 21N16 and 16N3 DNA are listed below (the sequence of Widom 601 DNA is underlined; the sticky ends are marked in bold):

21N16 DNA:

GATATCGAGCATCCGGATCCCCTGGAGAATCCCGGTGCCGAGGCCGCTCAATTGGTCGTAGACAGCTCTAGCACCGCTTAAACGCACGTACGCGCTGTCCCCCGCGTTTTAACCGCCAAGGGGATTACTCCCTAGTCTCCAGGCACGTGTCAGATATATACATCCTGTTCCAGTGCCGGTGCAC**CCC**;

16N3 DNA:

**CCC**GTGTCCAGTGCCGGTGCTGGAGAATCCCGGTGCCGAGGCCGCTCAATTGGTCGTAGACAGCTCTAGCACCGCTTAAACGCACGTACGCGCTGTCCCCCGCGTTTTAACCGCCAAGGGGATTACTCCCTAGTCTCCAGGCACGTGTCAGATATATACATCCTGTGAT

### 7. Reconstitution of histone octamers, nucleosomes, chromatosomes and asymmetric di-nucleosome

Histone octamers and nucleosomes were reconstituted by a previously published protocol.[4] Chromatosomes containing unmodified, monoubiquitylated or K63-linked-polyubiquitylated H1.0 were assembled as previously described with minor modifications.[5] Briefly, core histone octamers and 45N45 DNA were mixed at an equal stoichiometry in refolding buffer (10 mM Tris, pH 7.5, 2 M NaCl, 1 mM DTT). The salt concentration was gradient reduced from 2 M to 600 mM by using a peristaltic pump to add HE buffer (10 mM HEPES, pH 7.5, 1 mM EDTA). Next, unmodified, monoubiquitylated or K63-linked-polyubiquitylated H1.0 dissolved in 10 mM HEPES, pH 7.5, 600 mM NaCl, and 1 mM EDTA were added to the mixture. After additional dialysis for 2 h in 0.6 M NaCl buffer, the salt concentration was further gradient reduced from 600 to 200 mM. The mixture was finally dialyzed against HE Buffer for 8 h and purified by a Superose 6 Increase 5/150 GL column (GE Healthcare) pre-equilibrated in HE Buffer. Aliquotes from SEC were analyzed by native-PAGE. Aliquotes containing nucleosomes or chromatosomes were combined, concentrated to 2~3 μM, stored at 4°C, and used in one week.

To obtain asymmetric di-nucleosome, 21N16 and 16N3 DNAs were assembled into nucleosomes with Cy5–H2A, Fluo–H2A, H2B, H3, and H4. When gradient dialyzed against HE Buffer (10 mM HEPES, 1 mM EDTA, pH 7.5) till 600 mM NaCl, the former was supplemented with equimolar H1.0–K63Ub$_3$. After equilibrium dialysis for two hours, gradual dialysis was resumed to reduce salt concentration below 200 mM and then nucleosomes were transferred to another buffer (10 mM HEPES, pH 7.5, 1 mM DTT) for overnight. NanoDrop-Protein Label was used in Cy5 and Alexa 488 modes to quantify nucleosome concentrations, followed by mixing two types of nucleosomes at equimolar ratios in T4 DNA Ligase Buffer diluted to 2 × and adding T4 Ligase (LABLEAD, T5205) at a ratio of 1000 μg DNA per 1 unit, and transfer to pre-chilled Buffer (10 mM HEPES, pH 7.5, 1 mM DTT) for dialysis overnight. The reconstitution of asymmetric di-nucleosome was confirmed by 4.5% native PAGE imaging at Cy5 and Fluorescein mode (in red and green channel) respectively.

### 8. *In vitro* ubiquitylation assays

#### *8.1 Ubiquitylation assays on nucleosome, H1.0 chromatosome and H1.0–K63-Ub$_n$ chromatosomes*

Ubiquitylation assays on nucleosome, H1.0 chromatosome and H1.0–K63-Ub$_n$ chromatosome were performed in a reaction mixture containing 0.2 μM hUBA1, 1.0 μM UbcH5c, 2.0 μM RNF168, 100 μM Ub and 0.2 μM NCP or chromatosome in Ub assay buffer (50 mM Tris, 150mM NaCl, pH=7.6). Different RNF168 constructs and different NCP or chromatosomes were used according to related experimental purposes. H2A in NCP and chromatosomes was labeled with fluorescein at K129C for further analysis. Reactions were conducted at 37°C. Samples were taken at 0, 5, 10min and quenched by adding 4 × LDS loading buffer with 100mM DTT followed by boiling at 95°C for 5 min. Samples were resolved by SDS-PAGE on 12% NuPAGE Bis-Tris (Thermo Fisher Scientific) gels. The quantification of RNF168 activity was performed by calculating fluorescence intensity of each band using Image Lab 6.0.1. All reactions were performed in three independent biological replicates.

To analyze the contribution of different domains on RNF168 to activation effect on different templates, 2.0 μM RNF168$^{FL}$/ RNF168$^{1–193}$/ RNF168$^{1–159}$/ RNF168$^{1–113}$ and 0.2 μM 45N45 NCP/ H1.0 chromatosome/ H1.0–K63-Ub$_3$ chromatosome were used.

To analyze the impact of the length of ubiquitin chain on H1.0 on RNF168 activity, 2.0 μM $RNF168^{1\text{-}193}$ and 0.2 μM H1.0 chromatosome/ H1.0–Ub chromatosome/ H1.0–K63-$Ub_2$ chromatosome and H1.0–K63-$Ub_3$ chromatosome were used.

#### *8.2 Enzymatic Kinetics*

To perform the enzymatic kinetics assays, a series of $RNF168^{1\text{–}193}$ with a gradient of concentration was prepared. The reaction mixture contained 0.2 μM hUBA1, 1.0 μM UbcH5c, 100 μM Ub and 0.2 μM H1.0 chromatosome or H1.0–K63-$Ub_2$ chromatosome or H1.0–K63-$Ub_3$ chromatosome in Ub assay buffer (50 mM Tris, 150mM NaCl, pH 7.6). When the substrate was H1.0–K63-$Ub_2$ chromatosome or H1.0–K63-$Ub_3$ chromatosome, the gradient of $RNF168^{1\text{–}193}$ concentration was 0.25 μM, 0.5 μM, 1 μM, 2 μM, 4 μM, 8 μM and 16 μM. When the substrate was H1.0 chromatosome, the gradient of $RNF168^{1\text{–}193}$ concentration was 1 μM, 2 μM, 3 μM, 4 μM, 6 μM, 8 μM and 12 μM. The reaction was initiated by the addition of Ubiquitin. Samples were quenched at 10 min by adding 4 × LDS loading buffer with 100 mM DTT followed by boiling at 95°C for 5 min. After SDS-PAGE and quantification, the conversion ratio of H2A was fitted to the Michaelis-Menten equation to estimate the apparent catalytic constant $K_m$ in GraphPad Prism v.9.5.1.

#### *8.3 Ubiquitylation assays on asymmetric di-nucleosome*

0.1 μM hUBA1, 0.5 μM UbcH5c, 0.5 μM $RNF168^{FL}$ or $RNF168^{1\text{–}193}$, 0.1 μM asymmetric di-nucleosome were mixed in 4 × reaction buffer (100 mM ATP, 40 mM $MgCl_2$, 20 mM TCEP, pH 7.5). The reaction was incubated at 37°C. Samples were taken at 0.5, 2, 5, 10, 20, 30 min and quenched by adding 5 μL to 3 μL LDS loading buffer (containing 10 mM DTT) followed by boiling at 95°C for 5 min. Samples were resolved by SDS-PAGE on 4-12% NuPAGE Bis-Tris (Thermo Fisher Scientific) gels. The quantification of RNF168 activity was performed by calculating fluorescence intensity of each band under Fluorescein and Cy5 channel using Image Lab 6.0.1. All reactions were performed in three independent biological replicates.

### 9. Cryo-EM sample preparation

For the RNF168/UbcH5c–Ub/H1.0–K63-$Ub_3$ chromatosome complex (**CUP3**), freshly purified $RNF168^{1\text{–}193}$ (3.8 μM, final concentration) was mixed with freshly prepared H1.0–K63-$Ub_3$ chromatosome (1.9 μM, final concentration) at a stoichiometric ratio of 2:1. The mixture was incubated in a REC buffer (20 mM HEPES, pH 7.5, and 30 mM NaCl) at 4°C for 20 min. Next, the mixture was added to an equal volume of a REC buffer supplemented with 0.15%(v/v) glutaraldehyde. Glutaraldehyde crosslinking was performed at 25°C water bath for 10 minutes, and immediately quenched by 100 mM Tris·HCl (final concentration). The mixture was then by a

Superose 6 5/150 GL size-exclusion column (GE Healthcare) pre-equilibrated in REC buffer. Peak fractions were pooled and concentrated to ~150 ng·μL$^{-1}$ for cryo-EM sample preparation.

#### 10. Cryo-EM data collection and processing

A total of 1,164 cryo-EM micrographs were collected on a 300 kV Titan Krios cryo-electron microscope equipped with a Gatan K3 direct detector and a GIF quantum energy filter at a pixel size of 1.074 Å. micrographs were recorded at a dose of 50 electrons for 32 frames with a dose rate of 2.56 $e^-$ per frame. RELION v3.1.0 was used for data processing.[6] 951,132 auto-picked particles were extracted at a pixel size of bin4 (4.296 Å per pixel). Then, several rounds of reference-free fast 2D classification were performed to remove contaminations. After several parallel rounds of 3D classifications, particles containing both H1 and RNF168/UbcH5c densities were re-extracted at a full pixel size of bin1 (1.074 Å per pixel). The global resolution of the **CUP3** complex was calculated using the Fourier shell correlation (FSC) 0.143 criteria.[7] Local resolutions were estimated using the ResMap-1.1.4.[8]

#### 11. Model building

The refined and postprocessing maps (*B*-factor at 0) were used for model building, the nucleosomal DNA of 5NL0, the core histone octamer and RNF168, UbcH5c and Ub$^{H2A}$ of 8X7I, the AlphoFold model (AF-P07305-F1-model_v4) of linker histone H1.0 were merged into the initial model. The crucial model was then adjusted with residue point mutation (e.g., H1.0-K82C), invisible residue deletion and manual adjustment in WinCoot (0.8.2).[9] The model was followed with auto-refinement in Phenix1.19.2.[10] After several rounds of manual inspection and adjustment, the final model was given and validated. The detailed statistic was shown in **Table S1**.

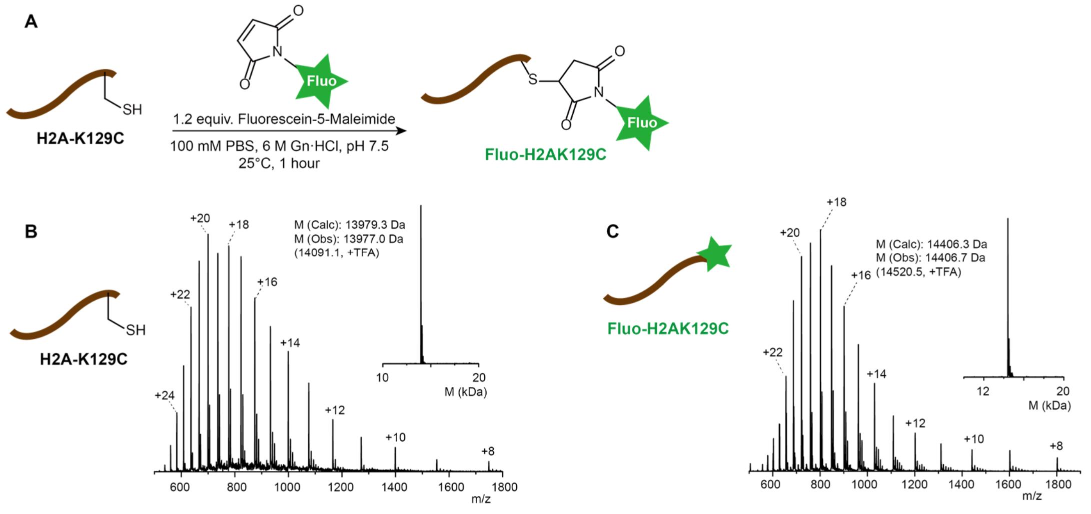


**Figure S1. Preparation and characterization of fluorescein-labeled H2A (Fluo–H2A).**

(A) Synthesis route diagram of Fluo–H2A.

(B) MS spectra and deconvoluted MS spectra of the starting material H2A-K129C.

(C) MS spectra and deconvoluted MS spectra of Fluo–H2A.

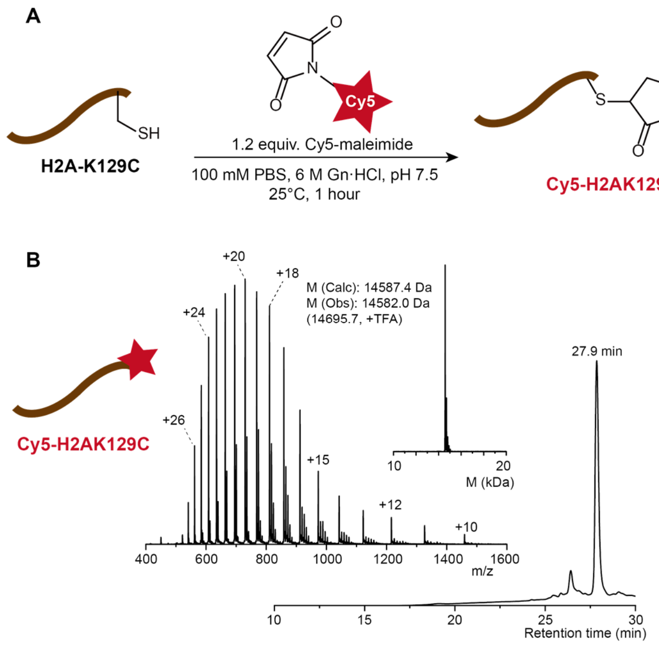


**Figure S2. Preparation and characterization of Cy5-labeled H2A (Cy5–H2A).**

**(**A) Synthesis route diagram of Cy5–H2A.

(B) RP-HPLC chromatograms (214 nm) and MS spectra and deconvoluted MS spectra of Cy5–H2A.

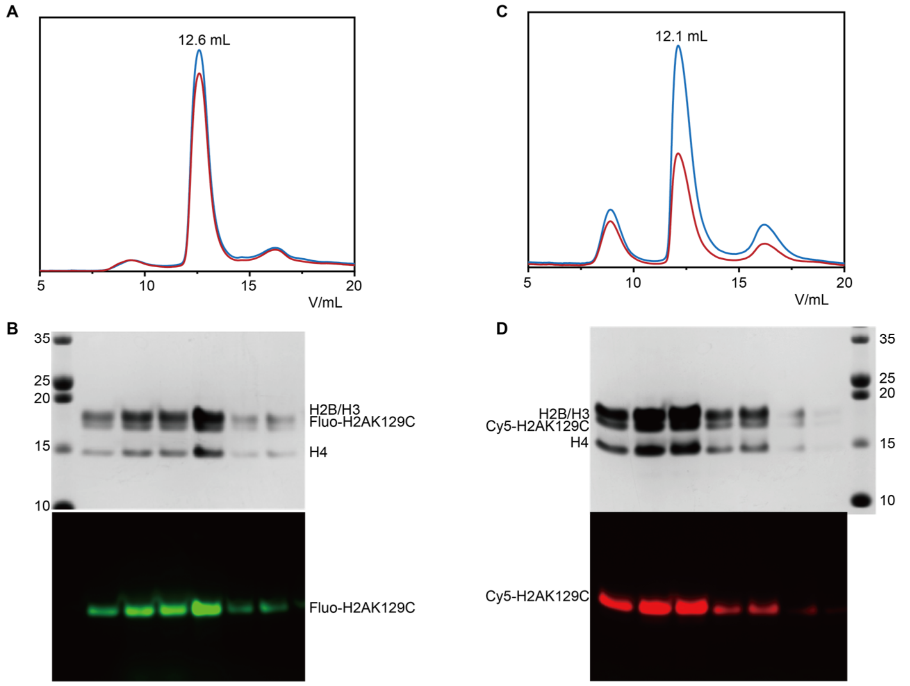


**Figure S3. Preparation of Fluo–H2A or Cy5–H2A histone octamer.**

(A) Size-exclusion chromatogram of Fluo–H2A histone octamer.

(B) SDS-PAGE gel images of Fluo–H2A histone octamer detected by Coomassie brilliant blue (CBB) staining (top) and Alexa 488 channel (green, bottom).

(C) Size-exclusion chromatogram of Cy5–H2A histone octamer.

(D) SDS-PAGE gel images of Cy5–H2A histone octamer detected by Coomassie-brilliant blue (CBB) staining (top) and Cy5 channel (red, bottom).

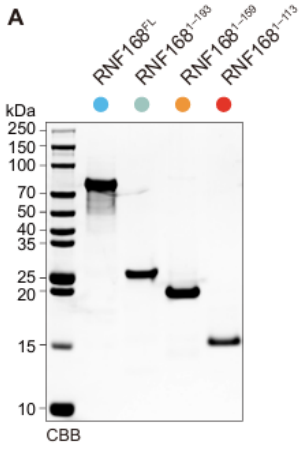


**Figure S4. Coomassie brilliant blue-stained SDS–PAGE gel of purified RNF168 constructs.**

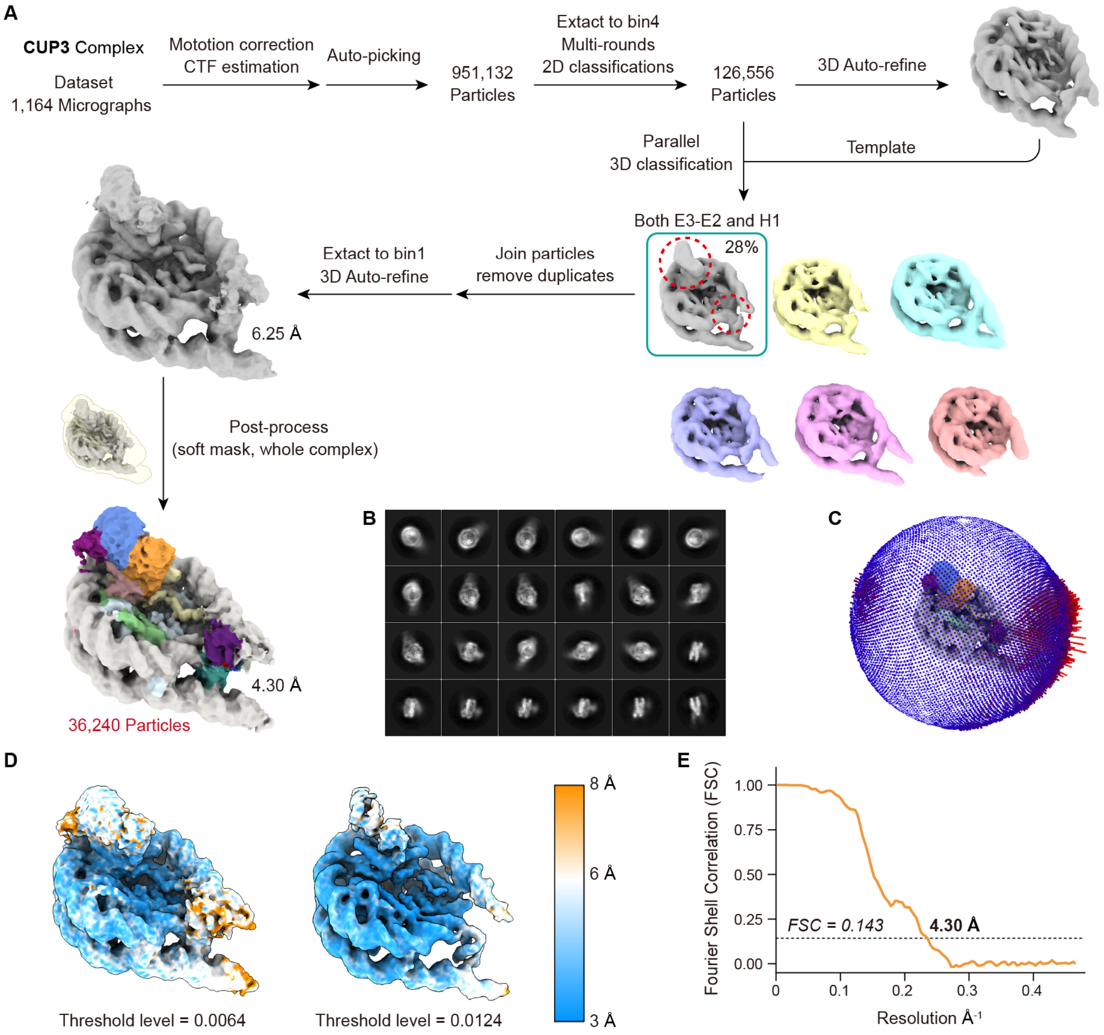


**Figure S5. Cryo-EM data processing for RNF168$^{1–193}$/Ubch5c–Ub/H1.0–K63-Ub$_3$ chromatosome (CUP3) complex.**

(A) Data processing flowcharts.

(B) Representative 2D classifications of the final reconstruction particles.

(C) Euler angle distributions of the final reconstruction particles.

(D) Local resolutions of the final cryo-EM reconstruction shown at the threshold levels of 0.0064 and 0.0124, respectively.

(E) Fourier shell correlation (FSC) curves for the final reconstruction.

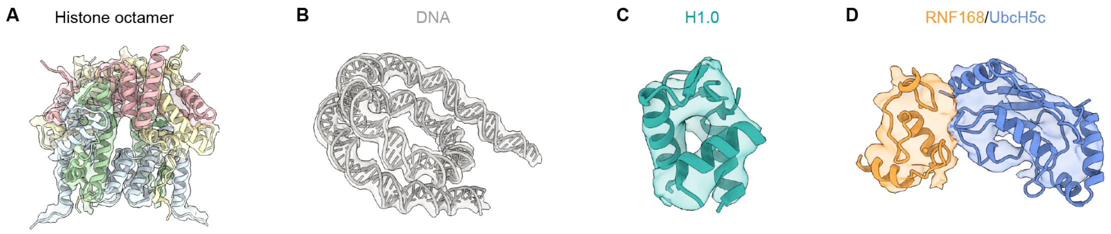


**Figure S6. Sample densities for CUP3 complex.**

(A) Histone octamer. (B) Nucleosomal DNA. (C) H1.0. (D) RNF168/UbcH5c module.

**Table S1. Cryo-EM data collection, refinement, and validation statistics**

| Data collection | RNF168$^{1\text{–}193}$/Ubch5c–Ub/H1.0–K63-Ub$_3$ chromatosome (CUP3) complex |
|---|---|
| PDB entry | 9IPU |
| EMDB entry | EMD-60781 |
| Voltage (keV) | 300 |
| Spherical aberration (mm) | 2.7 |
| Detector | K3 |
| Electron exposure | 50.0 $e^-$, 32 frames |
| Defocus range (μm) | -1.0 to -1.8 |
| Pixel size (Å) | 1.074 |
| Symmetry imposed | C1 |
| Box size (pixel) | 256 |
| Micrographs (no.) | 1,164 |
| Final particles (no.) | 36,240 |
| Map global resolution (Å) FSC threshold | 4.30 Å (0.143) |
| **Model Building and Refinement** | |
| Initial model used (PDB code) | 5NL0, 8X7I, AF-P07305-F1-model_v4 |
| Chains | 15 |
| Atoms | 15753 |
| Residues | Protein: 1130 Nucleotide: 340 |
| Ligands | two $Zn^{2+}$ |
| Cis proline/general (%) | 2.0/0.4 |
| Twisted prolin/general (%) | 0.0/0.0 |
| CaBLAM outliers (%) | 1.49 |
| Iso/Aniso (#) | 7662/8091 |
| Protein Min/max/mean | 30.00/306.25/130.33 |
| Nucleotide Min/max/mean | 257.66/299.01/278.34 |
| Bond lengths (Å) | 0.007 |
| Bond angles (°) | 1.110 |
| MolProbity score | 1.97 |
| Clashscore | 11.08 |
| Rotamer outliers (%) | 1.63 |
| Cβ outliers (%) | 0.00 |
| Favored (%) | 96.36 |
| Allowed (%) | 3.64 |
| outliers (%) | 0.00 |
